# The reovirus σ1 attachment protein influences the stability of its entry intermediate

**DOI:** 10.1101/2022.11.10.515810

**Authors:** Maximiliano L. Garcia, Pranav Danthi

**Affiliations:** Department of Biology, Indiana University, Bloomington, Indiana, USA

## Abstract

Structural metastability of viral capsids is pivotal for viruses to survive in harsh environments and to undergo timely conformational changes required for cell entry. Mammalian orthoreovirus (reovirus) is a model to study capsid metastability. Following initial disassembly of the reovirus particle mediated by proteases, a metastable intermediate called the infectious subvirion particle (ISVP) is generated. Using a σ1 monoreassortant virus, we recently showed that σ1 properties affect its encapsidation on particles and the metastability of ISVPs. How metastability is impacted by σ1 and whether the lower encapsidation level of σ1 is connected to this property is unknown. To define a correlation between encapsidation of σ1 and ISVP stability, we generated mutant viruses with single amino acid polymorphisms in σ1 or those that contain chimeric σ1 molecules composed of σ1 portions from Type 1 and Type 3 reovirus strains. We found that under most conditions where σ1 encapsidation on the particle was lower, ISVPs displayed lower stability. By separating wild-type particles based on the level of encapsidated σ1, we also found that lower σ1 encapsidation leads to lower ISVP stability. Characterization of mutant viruses selected for enhanced stability via a forward genetic approach also revealed that in some cases, σ1 properties influence stability without influencing σ1 encapsidation. These data indicate that σ1 can also influence ISVP stability independent of its level of incorporation. Together, our work reveals an underappreciated effect of the σ1 attachment protein on the properties of the reovirus capsid.

## INTRODUCTION

Proper attachment and cell entry are among the first steps for a successful virus infection. During the entry stage of the viral replication cycle, nonenveloped viruses undergo structural changes to allow for cytoplasmic delivery of the viral genomic material. A common strategy is for viruses to undergo conformational changes triggered by specific host factors or cellular environment, which in turn allow for the release of protein fragments or peptides that form pores in the endosome. For example, adenovirus is destabilized in the host endosome and releases protein VI, which disrupts the endosome (1–5). Poliovirus changes conformation upon receptor engagement, revealing VP4, which forms pores in the membrane (6–9). Maintaining proper temporal control of these conformational changes is critical for preserving viral infectivity. If virus conformation is altered prematurely, outside of the host, this could lead to an unsuccessful infection. Similarly, a virus that is too stable and fails to undergo conformational changes in a timely manner will become trapped in host compartments and never progress to the later steps of the replication cycle. Nonenveloped viruses maintain this balance through structural metastability of the capsid.

Mammalian orthoreovirus (reovirus) is a model for studying attachment and entry of non-enveloped viruses. Reovirus undergoes regulated conformational changes and its metastability influences efficiency of cell entry. Reovirus is comprised of two protein layers-the outer capsid and core, which contains a dsRNA genome. The particle binds junctional adhesion molecule A (JAM-A) and a glycan receptor via its σ1 protein (10, 11). σ1 is a trimer encapsidated in λ2 turrets and is a minor component of the outer capsid (12–15). Though a maximum of 12 σ1 trimers can be encapsidated on the viral particle, preparations of virus particles may contain particles with a range of σ1 trimers (16–18). After attachment, the virus enters the host cell through endocytosis, wherein the virion particle undergoes disassembly steps (19–23). The outermost capsid protein, σ3, is proteolytically degraded by acid-dependent proteases in endosomes (19). μ1, which resides underneath σ3, is then cleaved proteolytically at the C-terminus generating the Φ fragment which remains particle-bound (24–28). The degradation of σ3 and cleavage of μ1 forms infectious subvirion particles (ISVPs). ISVPs have an extended conformation of σ1 compared to virions (29). ISVPs convert to ISVP*s (22, 23, 27, 30–32). During this process, μ1 undergoes additional autocatalytic cleavage to generate μ1N. Further, Φ and μ1N are released and aid in membrane targeting and penetration (25–27). The σ1 trimer is also released upon conversion from ISVPs to ISVP*s (27, 30, 33).

Reovirus can undergo reassortment in cell culture and in vivo (34). Our recent work highlights unexpected phenotypes of reassortant strains that contain one or more capsid protein genes from a different parental strain. In one such study, we used a reassortant that expresses the σ1 protein of strain type 3 Dearing (T3D) Cashdollar (T3DC) in an otherwise T3D Fields (T3D^F^) virus (35, 36). Although the σ1 proteins of T3DC and T3D^F^ differ only by two amino acids, the reassortant virus (T3D^F^/T3D^C^S1) displays many phenotypes. The encapsidation of σ1 on particles T3D^F^/T3D^C^S1 is lower. Specifically, within purified preparations, a large proportion of T3D^F^/T3D^C^S1 virus particles contain either no σ1 trimers or a low number of encapsidated σ1 trimers. T3D^F^/T3D^C^S1 particles are less stable than those of T3D^F^, and consequently convert to ISVP*s more readily. T3D^F^/T3D^C^S1 produces smaller plaques and is also less infectious than T3D^F^ (36). Based on these results, we hypothesized that mismatched interaction between T3DC σ1 and the remainder of the T3D^F^ particle as a consequence of reassortment alters optimized protein-protein interactions within the particle and produces these effects. However, the precise basis for how these two amino acid changes in σ1 produced all of these phenotypes and how these phenotypes relate to each other remains unknown.

In this study, through the use of point mutants, we determined that each of the polymorphic T3D^F^ and T3DC σ1 residues, 22 or 408 of σ1, contribute to ISVP stability. We also found changes at each of these residues altered the encapsidation pattern of σ1 on the particle. This finding suggested a possible link between ISVP stability and σ1 encapsidation in certain contexts. To define whether σ1 had a function in ISVP stability held true in other contexts, viruses with chimeric σ1 proteins, comprised of σ1 domains swapped between two different reovirus strains, were used. Chimeric σ1 proteins are not incorporated into particles with efficiency similar to the cognate σ1 protein and ISVPs of these viruses also display lower stability. Finally, upon separating wild-type particles based on their σ1 levels, we observed that even for wild-type viruses, the levels of σ1 on the particle correlate with ISVP stability. Thus, incorporation levels of σ1 influence ISVP stability. Using forward genetics, we selected a revertant that restores the stability of ISVPs of T3D^F^/T3DCS1. Recovery of an engineered virus indicated that a single amino acid substitution in the distal portion of σ1 restores ISVP stability, but did not affect encapsidation of σ1. This result along with evidence that some viruses with chimeric σ1 molecules display lower stability without influencing σ1 encapsidation indicate that σ1 impacts ISVP stability via a second, distinct mechanism.

## RESULTS

### Residues in the σ1 head and tail domain impact ISVP stability

The σ1 proteins of T3D^F^ and T3D^C^ vary by only two amino acids – a valine-to-alanine polymorphism at residue 22 and threonine-to-alanine difference at residue 408 (35). To determine which of these residues impart phenotypes described for T3D^F^/T3D^C^S1, viruses with each individual change was constructed in a T3D^F^ background (T3D^F^/S1-22^C^ or T3D^F^/S1-408^C^). In vitro generated ISVPs were incubated at a range of temperatures, treated with trypsin, and resolved on SDS-PAGE gels. The native conformation of the μ1 δ fragment present on ISVPs is resistant to trypsin digestion. Trypsin digests δ at residues that are exposed due to conformation changes induced by heat. We used the disappearance of the δ band by trypsin treatment as an indicator of conversion of δ from a native to an altered, ISVP* conformer (Fig. 1A) (27, 37). Under the conditions used, T3D^F^ failed to convert to ISVP*s at temperatures up to 38°C. In contrast, consistent with previous work, T3D^F^/T3D^C^S1 converted to ISVP* at a significantly lower temperature of 36°C (36). Interestingly, we found that T3D^F^/S1-22^C^ and T3D^F^/S1-408^C^ also converted to ISVP*s at a lower temperature of 35-36°C. Thus, changes at either residue 22 or 408 in σ1 is sufficient to lower stability of T3D^F^ ISVPs.

**Figure 1.**
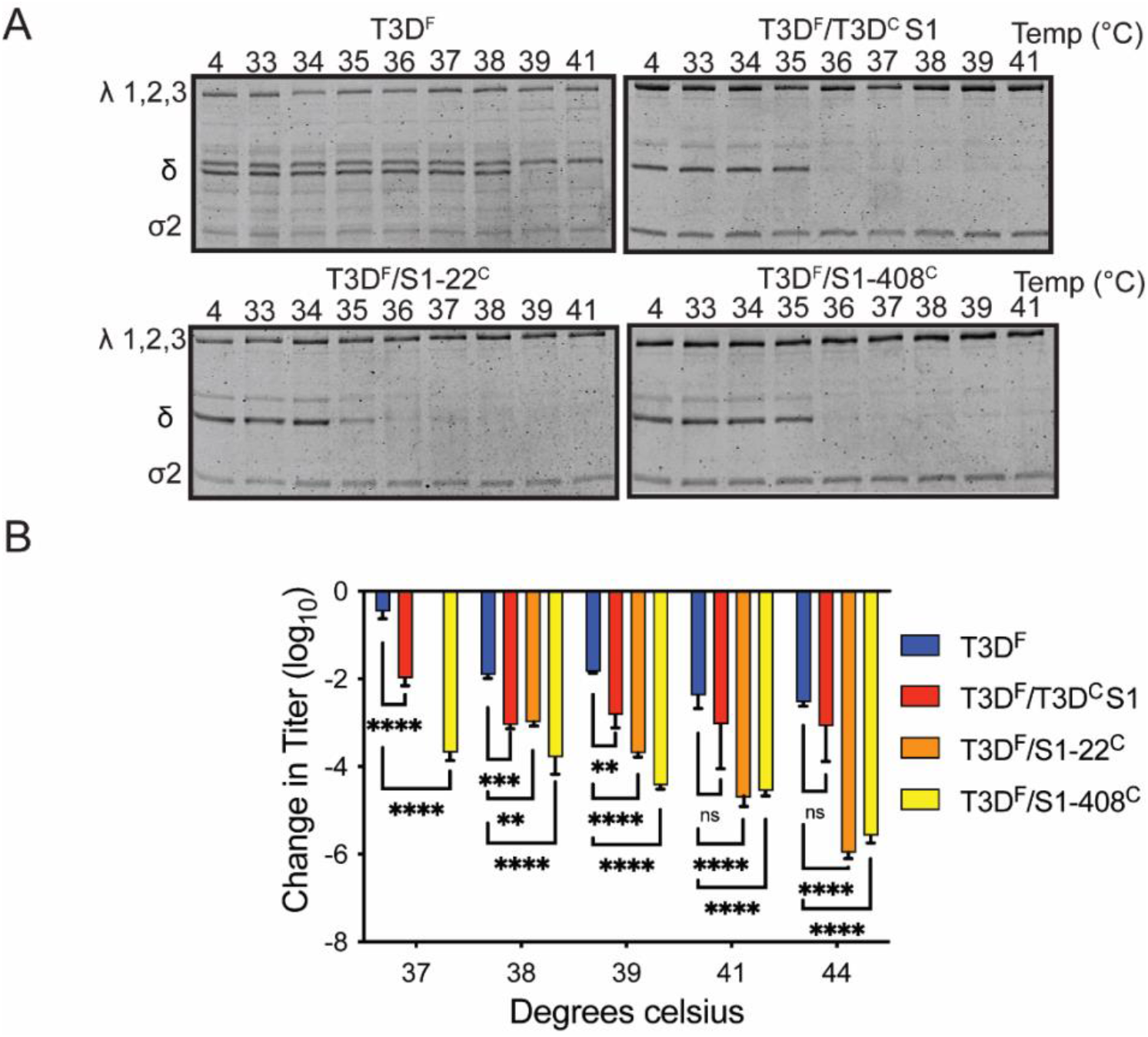
T3D^F^/T3DCS1, T3D^F^/S1-22C, and T3D^F^/S1-408C ISVPs are less stable than T3D^F^. (A) ISVPs (2×10^12^ particles/mL) of T3D^F^, T3D^F^/T3DCS1, T3D^F^/S1-22C, or T3D^F^/S1-408C were incubated at the indicated temperatures for 5 min. After heat treatment, the aliquots were incubated with 0.1 mg/ml trypsin for 30 min on ice. Loading dye was added and samples were heated at 95°C for 5 min. Samples were resolved on 10% SDS-PAGE gels. These gels represent at least 3 independent experiments. (B) ISVPs of T3D^F^, T3D^F^/T3D^C^S1, T3D^F^/S1-22^C^, or T3D^F^/S1-408^C^ were heated at a range of temperatures. After heating, the samples were subjected to plaque assay. Change in titer (represented as log10 PFU/mL) at each temperature relative to 4°C was measured. Data is representative of 3 independent experiments. Error bars represent SD of 3 replicates. Two-way ANOVA with Dunnett’s multiple comparisons was performed. **, ***, **** represents, *P* < 0.001,0.0005, 0.0001 respectively.

In vitro conversion of ISVP to ISVP*s results in a decrease in viral titer (27, 33, 38). As an additional measure for quantifying the stability of ISVPs, particles incubated at a range of temperatures were subjected to a plaque assay (Fig. 1B). In comparison to ISVPs of T3D^F^, ISVPs of T3D^F^/T3D^C^S1 lose titer to a greater extent at temperatures below 39°C, consistent with their lower stability. At 41 °C or higher, the loss of titer for ISVPs of T3D^F^ and T3D^F^/T3D^C^S1 is similar. T3D^F^/S1-22^C^ and T3D^F^/S1-408^C^ both displayed significantly greater loss in titer compared to T3D^F^ at nearly every temperature. These data confirm that either a change at residue 22 or at 408, is sufficient to compromise the stability of T3D^F^ ISVPs.

In comparison to T3D^F^, T3D^F^/T3D^C^S1 encapsidates a lower level of σ1 on particles (36). To determine if the changes at residues 22 and 408 impact the amount of σ1 encapsidated into the particle, we coated each of these types of particles on to plates and measured the encapsidation of σ1 relative to the major outer-capsid proteins σ3 and μ1 using FLISA. Consistent with previous work, T3D^F^ contained significantly more σ1 than T3D^F^/T3D^C^S1 (Fig. 2A) (36). We found that the amount of σ1 was lower on both T3D^F^/S1-22^C^ and T3D^F^/S1-408^C^, though only T3D^F^/S1-408^C^ showed a statistical difference. Reovirus particles encapsidate a range of 0-12 σ1 trimers on particles (17, 18). Resolving purified particles on native agarose gels results in separation of these populations based on σ1 content (Fig. 2B, 2C) (18). Using this technique, we confirmed our previous result that purified T3D^F^ preparations contain more particles with higher levels of encapsidated σ1 compared to T3D^F^/T3D^C^S1(36). We found that T3D^F^/S1-408^C^ particles resemble T3D^F^/T3D^C^S1 in the amount of encapsidated σ1. In the preparation of T3D^F^/S1-22^C^, though the number of particles with very few (0-3) trimers resemble T3D^F^, there appear to be fewer particles with > 4 σ1 trimers. Thus, our data indicate that although residue 22 only has a modest effect on total amount of σ1 encapsidated in a preparation of particles, it does change the fraction of particles in a population that contain high and low level of σ1 incorporation. The altered σ1 encapsidation pattern of T3D^F^/T3D^C^S1, T3D^F^/S1-22^C^ and T3D^F^/S1-408^C^ may be related to the lower expression of lower stability of mutant σ1 in infected cells. To evaluate this possibility, σ1 levels in infected cells were measured using western blot (Fig. 2D). We found equivalent levels of σ1 following infection with each of the four viruses, implying that the deficiency in σ1 encapsidation is not from insufficient production or accumulation of σ1. Thus, these data suggest that changes to residue 22 and 408 influence the capacity of σ1 to become encapsidated or remain encapsidated.

**Figure 2.**
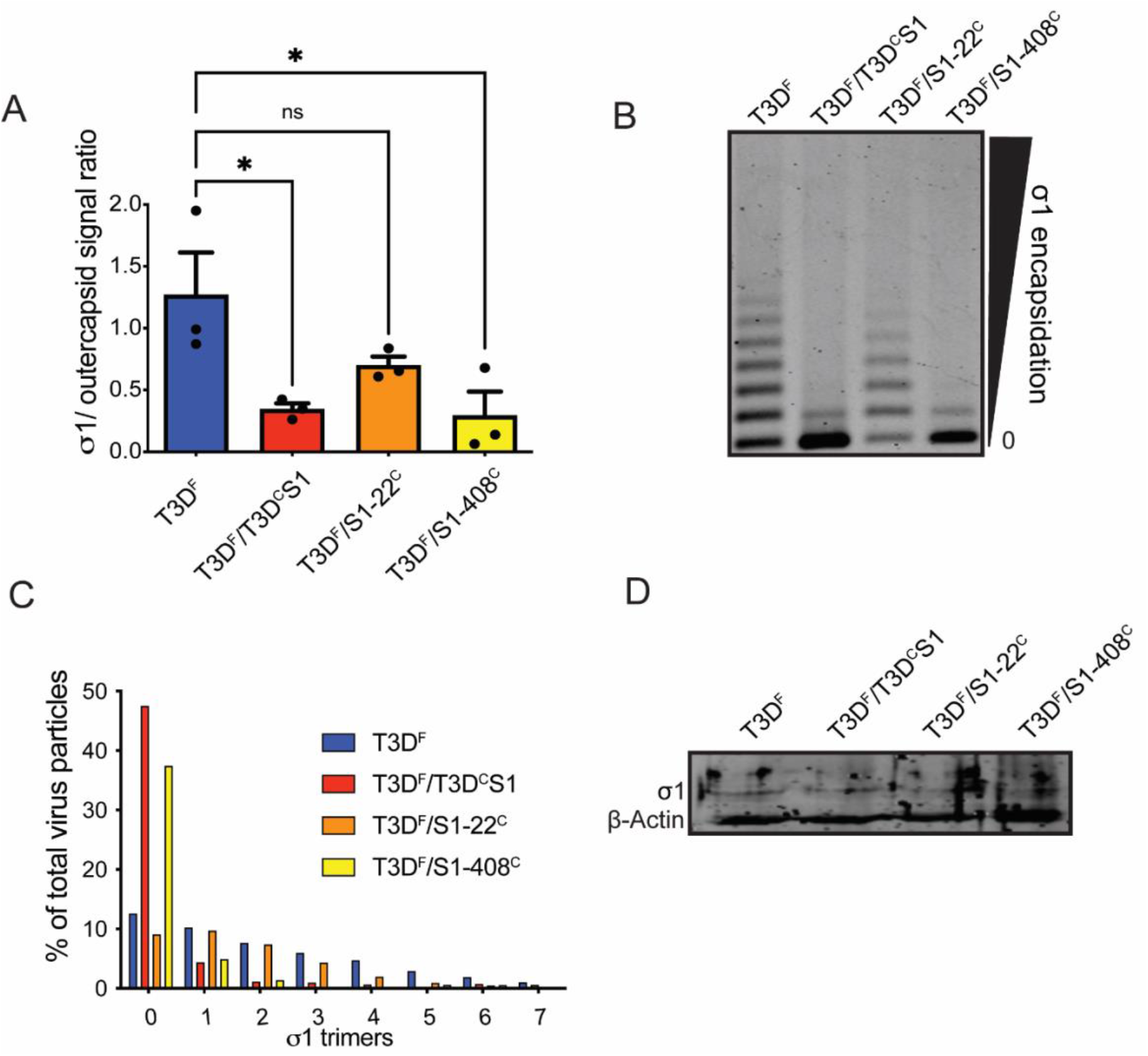
Residues in the head and tail domains of T3D σ1 impact stability and encapsidation. (A) Purified T3D^F^, T3D^F^/T3D^C^S1, T3D^F^/S1-22^C^, or T3D^F^/S1-408^C^ virions were diluted in 0.1M carbonate-bicarbonate buffer and coated overnight on to high-affinity-binding polystyrene plates at 4°C. After blocking, the virus was stained with reovirus specific rabbit polyclonal antiserum or σ1 specific 9BG5 mAb. After labeling with appropriate secondary antibodies the plate was imaged on LI-COR Odyssey scanner. The results are plotted as the ratio of 9BG5 intensity compared to reovirus polyclonal intensity in the same well. 3 wells were counted and error bars represent SD. *, *P*<0.05 as determined by Students *t* test compared to T3D^F^ (B) 1×10^11^ virions were resolved on an ultra-pure agarose gel based on their σ1 trimer content. After colloidal blue staining, the gel was imaged on a LI-COR Odyssey scanner. (C) The percent of particles present in each species was quantified relative to the total intensity of one lane. (D) L929 cells were infected by T3D^F^, T3D^F^/T3D^C^S1, T3D^F^/S1-22^C^, or T3D^F^/S1-408C at an MOI of 5. Lysates were collected and subjected to western blot. Antibodies against T3D σ1 head and β-Actin were used. Membranes were scanned on a LI-COR Odyssey scanner to visualize σ1 and β-Actin bands.

### ISVPs of viruses encapsidating T1L-T3D chimeric σ1 molecules display lower stability

While the studies presented above suggest a link between the pattern or level of σ1 encapsidation and ISVP stability in the T3D^F^ genetic background, different reovirus strains often display strain-specific phenotypes. To determine if ISVP stability is similarly affected by σ1 in other genetic backgrounds, we used previously characterized T1L viruses that express chimeric σ1 proteins from T1 and T3 reoviruses (39). We reasoned that if these viruses with chimeric σ1 proteins display lower levels of ISVP stability, this would strengthen the connection between ISVP stability and σ1. Further, because T1 and T3 σ1 proteins display substantial level of amino acid differences despite their structural similarity, these viruses could also give insight into which domains of σ1 affect these phenotypes (15, 40, 41). The σ1 protein can be divided into the tail, body, and head domain (Fig. 3A) (41). We used viruses that express a σ1 chimera containing only the T3 tail domain (T3 tail) and a σ1 chimera expressing the T3 tail and body domain (T3 tail+T3 body) (39). In comparison to T1L, T3 tail and T3 tail+T3 body display less stable ISVPs (Fig. 3B). While T1L ISVPs convert to ISVP*s at 55°C, ISVPs of T3 Tail and T3 tail+T3 body convert to ISVP*s at 52°C. These results provide additional evidence that σ1 properties impact ISVP stability.

T1L, T3 tail, and T3 tail +T3 body were also assayed to determine both the total encapsidation, and the pattern of σ1 encapsidation (Fig. 4A). Using a combination T1 σ1-specific and outer-capsid specific antisera and FLISA assays we determined that all three viruses displayed an equivalent level of total σ1, consistent with previous work (39). Using separation on native agarose gels, we found that T1L preparation contains particles with a range of encapsidated σ1 (Fig. 4B, 4C). Yet, the preparation predominantly contains particles with high level of σ1 encapsidation. T3 Tail preparations primarily display particles with highly encapsidated σ1 trimers but also contain some particles with lower level σ1 encapsidation (a similar population is not detectable in T1L). T3 tail+ T3 body preparations have the majority of their particles encapsidated with 0 to 3 σ1 trimers. Thus, despite their similarity to T1L in the total amount of encapsidated σ1, T3 Tail and T3 tail+T3 body viruses possess a population of virions with little to no σ1. Much like our data with changes at residue T3D σ1 residue 22 above, these data suggest ISVP stability may not be simply linked to the total amount of encapsidated σ1 but rather to the proportion of particles that contain a lower or higher level of encapsidation. It is also possible that the effect of σ1 on ISVP stability in these contexts is not related to the amount of encapsidation.

**Figure 3.**
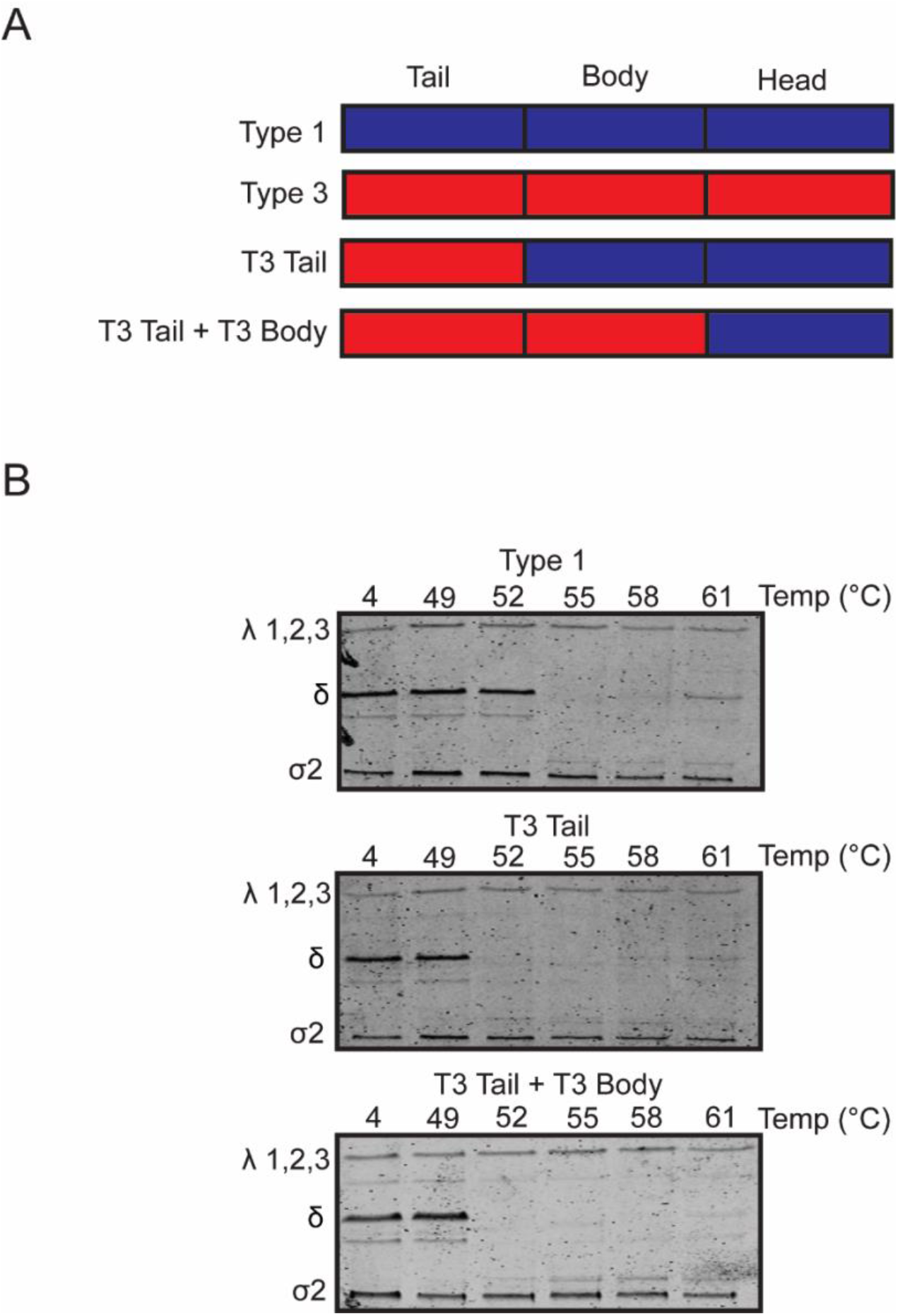
Tail and body domains of σ1 influence ISVP stability. (A) schematic representation of type 1 virus (T1), type 3 virus (T3) and chimeric σ1 proteins. (B) ISVPs (2×10^12^ particles/mL) of T1L, T3 Tail, or T3 Tail + T3 Body were incubated at the indicated temperatures for 5 min. After heat treatment, the aliquots were incubated with 0.1 mg/ml trypsin for 30 min on ice. Loading dye was added and samples were heated at 95°C for 5 min. Samples were resolved on 10% SDS-PAGE gels. These gels represent at least 3 independent experiments

**Figure 4.**
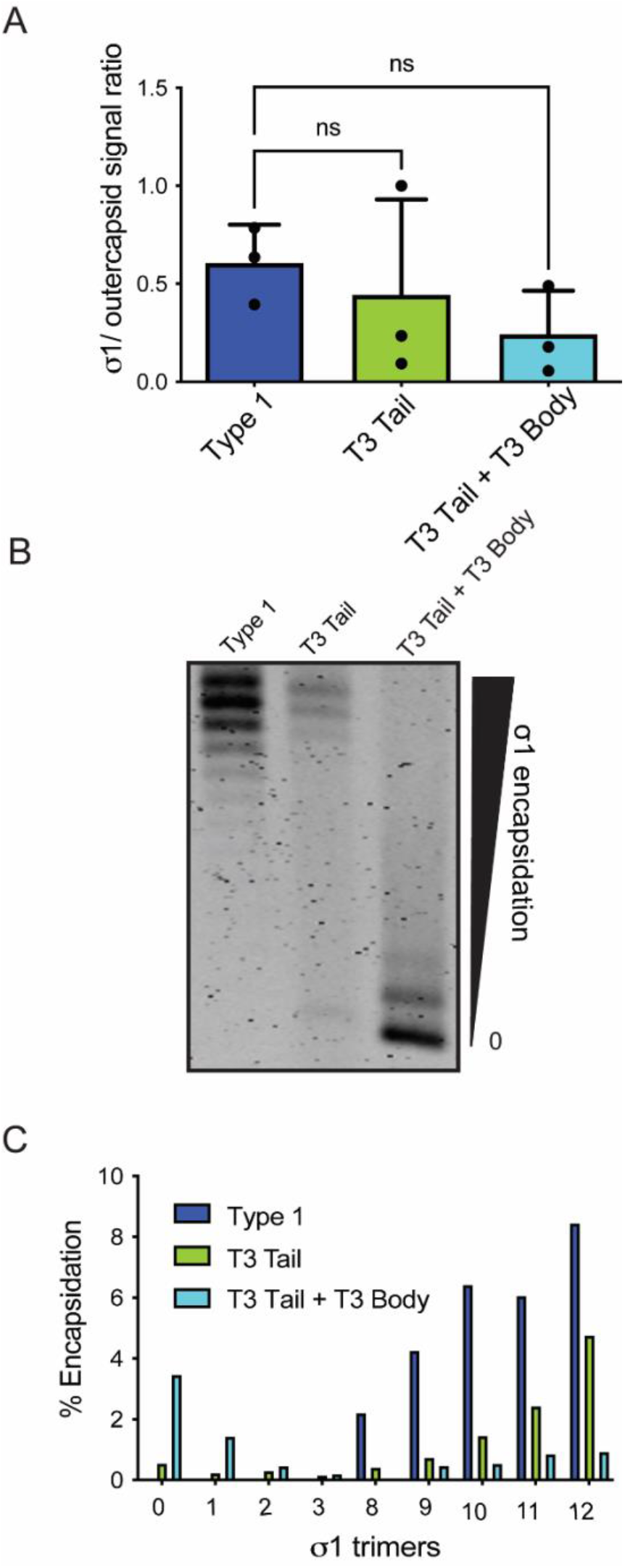
Alterations in σ1 tail and body domains affect its encapsidation on particles. (A) Purified T1L, T3 Tail, or T3 Tail+T3 Body virions were diluted in 0.1M carbonate-bicarbonate buffer and coated overnight on to high-affinity-binding polystyrene plates at 4°C. After blocking, the virus was stained with reovirus specific rabbit polyclonal antiserum or σ1 specific 5C6 mAb. After labeling with appropriate secondary antibodies the plate was imaged on LI-COR Odyssey scanner. The results are plotted as the ratio of 5C6 intensity compared to reovirus polyclonal intensity in the same well. 3 wells were counted and error bars represent SD. *, *P* <0.05 as determined by Students *t* test compared to T1L. (B) 1×10^11^ virions were resolved on an ultra-pure agarose gel based on their σ1 trimer content. After colloidal blue staining, the gel was imaged on LI-COR Odyssey scanner. (C) The percent of particles in each trimer species was quantified relative to the total.

### Wild-type ISVPs with lower σ1 encapsidation are less stable

Our experiments above suggest a link between σ1 encapsidation patterns and ISVP stability in some contexts. However, since all of our experiments used mutant viruses, we cannot rule out the possibility that σ1 encapsidation and ISVP stability are impacted independently by the same mutation. To address this, we sought to separate groups of wild type viruses based on level of σ1 encapsidation. Toward this goal, T1L was resolved on native agarose gels in multiple lanes. One lane was stained with ethidium bromide to ascertain the positions of virions containing different amounts of σ1. The other lanes were cut into three parts such that particles with varying σ1 encapsidation (low, medium, and high) would be present in different groups (Fig. 5A). The virus within these gel portions was extracted, converted to ISVPs, and subjected to incubation at various temperatures and its stability was measured by plaque assay. Particles with low σ1 remained infectious as ISVPs at 4°C, but upon incubation at higher temperatures, converted to ISVP*s and lost all infectivity. Particles with a moderate level of encapsidated σ1 shared the same ISVP stability as particles with low σ1 encapsidation and converted to ISVP*s. However, particles with high amounts of σ1 maintained ISVP stability up to 45°C (Fig. 5B). These data indicate that even for viruses with identical genotypes (in this case, wild-type) ISVPs with a higher level of encapsidated σ1 have greater thermostability. Thus, one way in which σ1 properties impart stability on the particle is via encapsidation.

**Figure 5.**
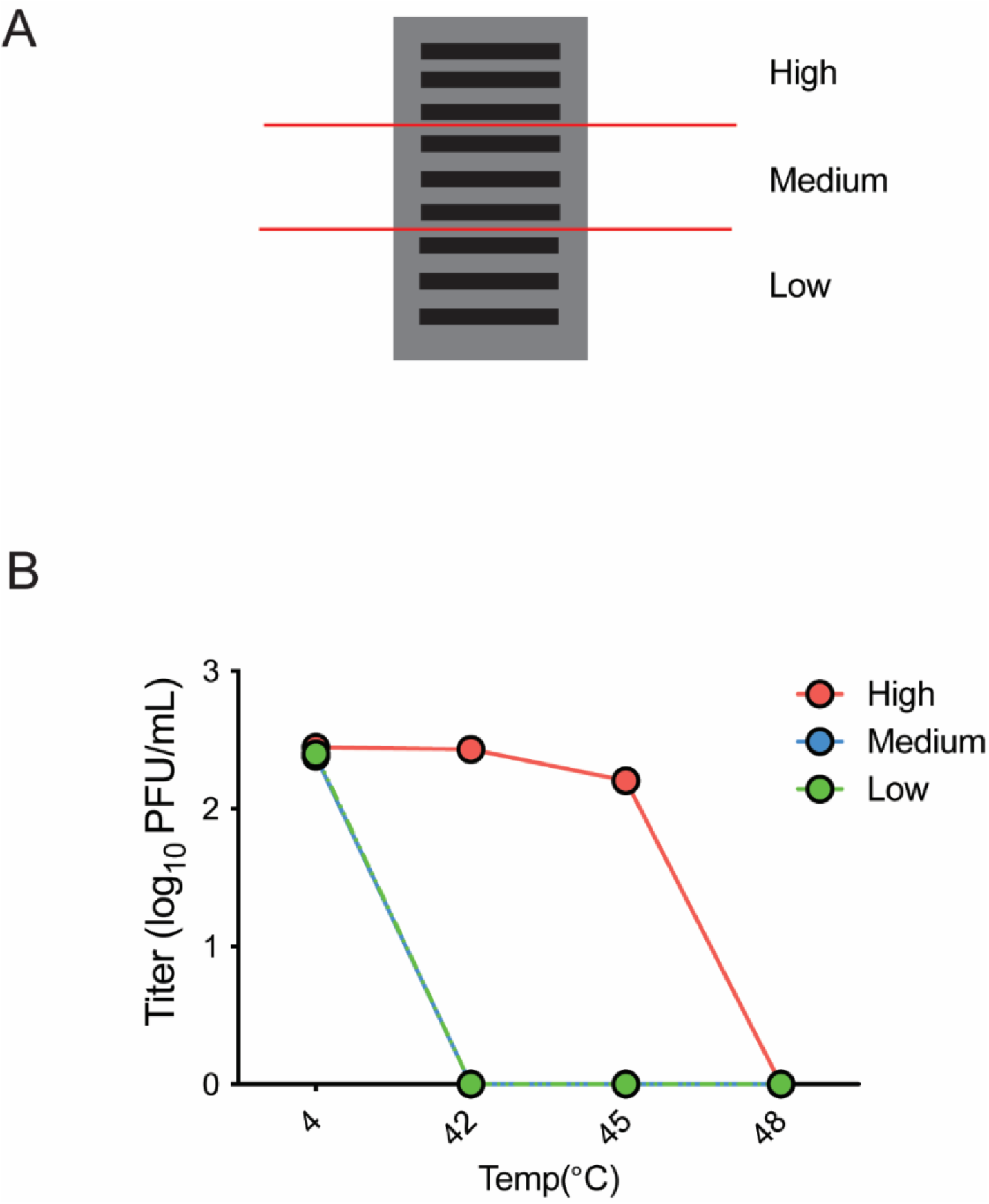
σ1 encapsidation level of wild-type virus affects ISVP stability. (A) Diagram of T1L separated based on encapsidation status. (B) 1×10^11^ virions were resolved on an ultra-pure agarose gel based on their σ1 trimer content. The same sample was run on multiple lanes. One lane was cut and stained for 3 hours with ethidium bromide. Using the stained lane as guide, the other lanes were cut into groups based on the amount of encapsidated σ1. Samples were minced, sonicated, and nutated. The released virus was converted to ISVPs and heated at a range of temperatures. After heating, the samples were subjected to plaque assay. Change in titer (represented as log_10_ PFU/mL) at each temperature relative to 4°C was measured. Data is representative of 3 independent experiments.

### Selection of thermostable revertants reveals a stabilizing mutation in σ1

To gain more information about how σ1 affects ISVP stability, we performed a forward genetic screen to isolate stabilizing revertants of T3D^F^/T3D^C^S1. T3D^F^/T3D^C^S1 was converted to ISVPs and heated at a temperature that typically destabilizes the particles and results in a loss of infectivity. Heat resistant (HR) variants within this population that survived this treatment and formed plaques were deemed to have changes that stabilized the particles. These plaques were picked, and the thermostability of ISVPs of these plaque isolates was confirmed (Fig. 6A). The S1, L2 and M2 genome segments which respectively encode σ1, λ2 and μ1 of these viruses were sequenced. These genome segments were selected for sequencing because σ1 is the subject of this study and σ1 interacts with λ2 (12–15). Further, μ1 is a known regulator of ISVP stability (33, 38, 42, 43). Point mutations were found at residue 132 in σ1, at residues 857 and 551 in λ2, and residue 594 in μ1. From these, the basis for the stability of HR5 was found to be most interesting because it had acquired a threonine-to-alanine mutation in σ1 at residue 132 offering us an opportunity to further understand how σ1 could impact ISVP stability. In the structure of recombinantly expressed σ1, residue 132 resides in the trimerization domain (Fig. 6B). However, this residue is not expected to be in a position where it could interact with other monomers of σ1 (Fig. 6C).

**Figure 6.**
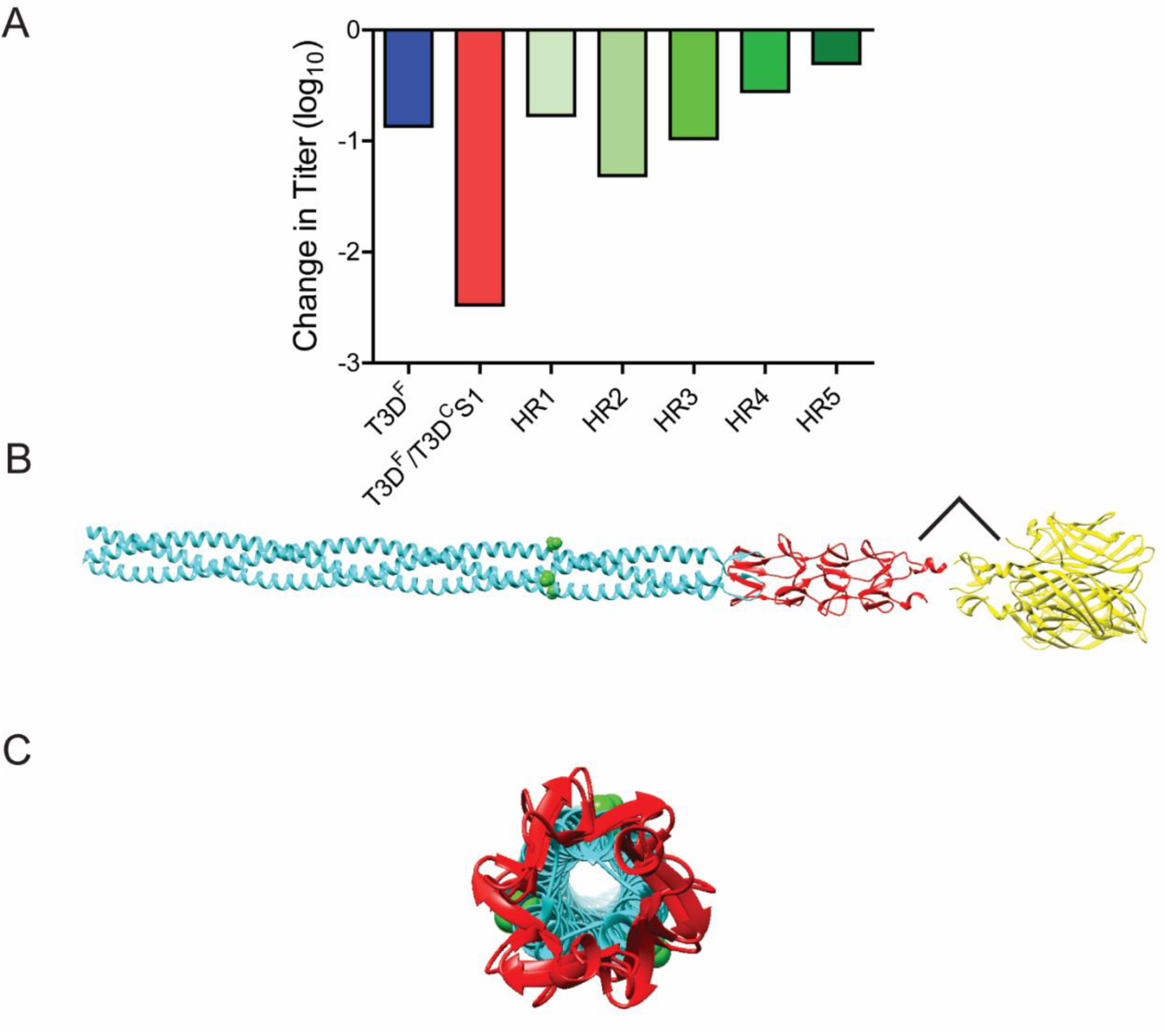
Selection of thermostable revertant, T3D^F^/T3D^C^S1-T132A. (A) Candidate thermostable revertants were converted to ISVPs and incubated at 4°C or 44°C. After incubation the samples were subjected to plaque assay to determine the titer. Change in titer (represented as log10 PFU/mL) at 44°C relative to 4°C was measured. (B, C) UCSF Chimera rendered image of a side view (B) or top view (C) of the σ1 trimer. PDB accession number (6GAP) (2OJ5). The position of the mutated threonine at residue 132 is highlighted by green spheres.

### T3D^F^/T3D^C^S1-T132A partially restores ISVP stability, but not σ1 encapsidation

To isolate the role of this σ1 change from the remainder of the genotype of HR5, we generated T3D^F^/T3D^C^S1-T132A via reverse genetics. This strain was purified and compared to T3D^F^/T3D^C^S1. To determine if T132A impacts ISVP stability, the temperature at which the δ fragment acquires a protease-sensitive conformation was determined (Fig. 7A). Based on the protease sensitivity of their δ fragments, T3D^F^/T3D^C^S1 converted to ISVP*s at 38°C, while T3D^F^/T3D^C^S1-T132A converted to ISVP*s at 41 °C. Since the parental virus, T3D^F^ converts to ISVP*s at 44°C (Fig. 1), these data indicate that T132A in σ1 partially restores the stability of T3D^F^/T3D^C^S1. To determine if T132A change in σ1 restores ISVP stability by fixing the encapsidation defect in T3D^F^/T3D^C^S1, we compared both the total level of σ1 and the pattern of σ1 incorporation between T3D^F^/T3D^C^S1 and T3D^F^/T3D^C^S1-T132A. FLISA assays demonstrated that the encapsidation of σ1 was not significantly increased in T3D^F^/T3D^C^S1-T132A compared to T3D^F^/T3D^C^S1 (Fig. 7B). Similarly, particles with little to no σ1 were equally abundant in T3D^F^/T3D^C^S1-T132A and T3D^F^/T3D^C^S1 (Fig. 7C, E). This demonstrates that σ1 can also affect ISVP stability even when encapsidation of σ1 remains low.

**Figure 7.**
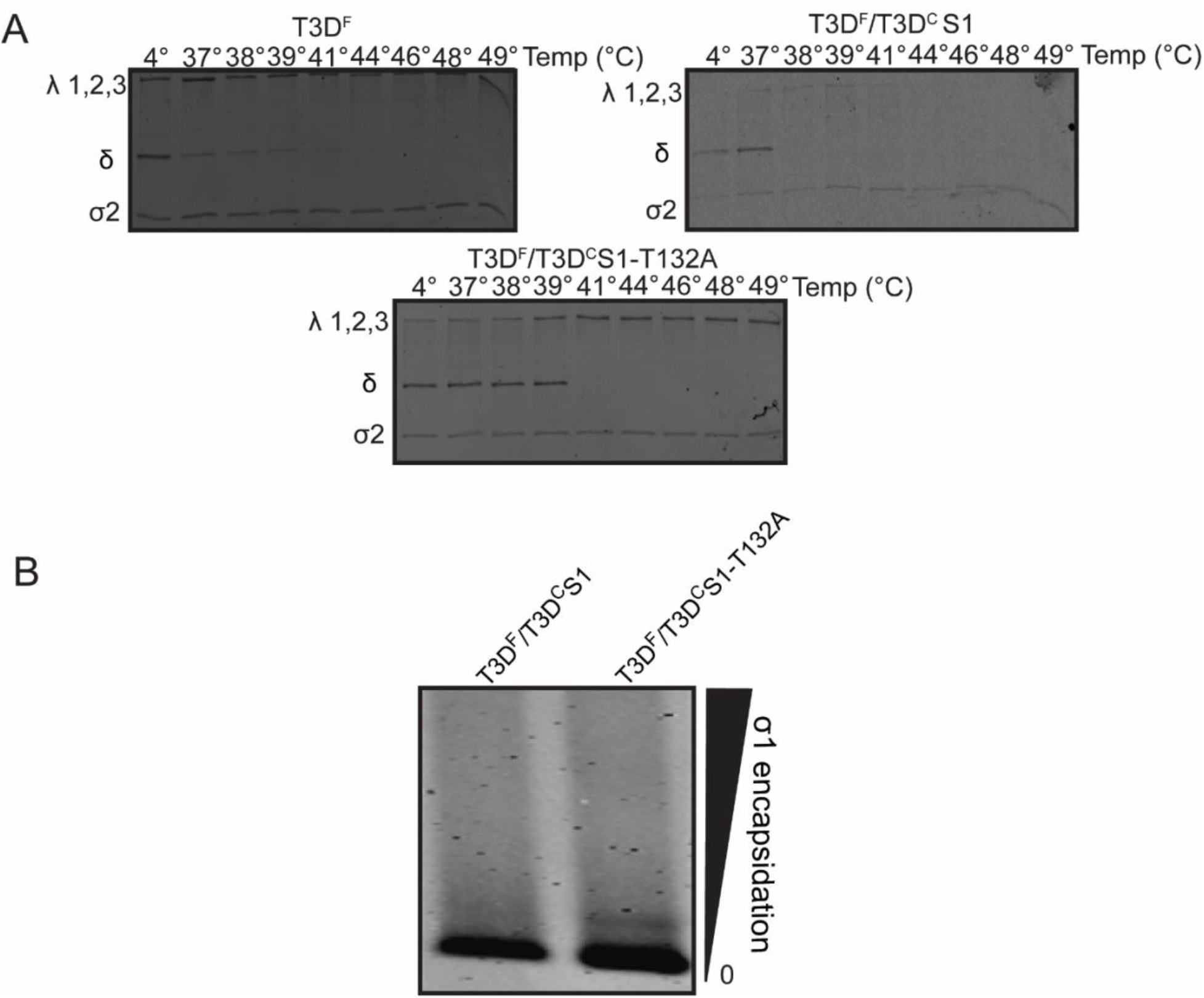
σ1 of T132A restores T3D^F^/T3D^C^S1 ISVP stability and enhances encapsidation of σ1. (A) ISVPs (2×10^12^ particles/ml) of T3D^F^, T3D^F^/T3D^C^S1, or T3D^F^/T3D^C^S1-T132A were incubated at the indicated temperatures for 5 min. After heat treatment, the aliquots were incubated with 0.1 mg/ml trypsin for 30 min on ice. Loading dye was added and samples were heated at 95°C for 5 min. Samples were run on SDS-PAGE gels. These gels represent at least 3 independent experiments. (B) 1×10^11^ virions were resolved on an ultra-pure agarose gel based on their σ1 trimer content. After colloidal blue staining, the gel was imaged on LI-COR Odyssey scanner.

## DISCUSSION

Our previous work using T3D^F^ and T3D^F^/T3D^C^S1 revealed an unexpected link between the properties of σ1 and the stability of ISVPs (36). Here we sought to further understand this link. We found that changes in σ1 that reduce its encapsidation on particles, diminish ISVP stability (Fig 2). Wild-type particles separated on the basis of their σ1 levels also demonstrated that particles with lower σ1 have lower ISVP stability (Fig 5). Based on these data we conclude that the presence of σ1 on the particle directly affects ISVPs stability. Our work also revealed that for some σ1 mutant viruses σ1 encapsidation levels and ISVP stability do not correlate. These data indicate that σ1 may impact ISVP stability in more than one way. Together our work provides new information to increase an understanding of how the attachment fiber of reovirus impacts the stability of ISVPs.

σ1 is comprised of 3 structural domains. The globular head, which is responsible for binding JAM-A, β-sheet repeats, which form the body and bind secondary glycans for type 3 reoviruses, and finally the coiled coil, which forms the tail forming trimers and anchoring σ1 into λ2 (10, 11, 44, 45). The function of σ1 in mediating attachment to cell surface receptors is well characterized but information about its role in controlling particle stability is lacking (10, 11, 46–50). Our work indicates that alterations in single residues in the tail or head and entire domains in the tail or body decrease encapsidation, demonstrating that all domains play some role in encapsidation, but to varying extents. It is not surprising that altering the tail would affect encapsidation. The importance of the hydrophobic tail of σ1 in anchoring into λ2 has been described (45). Deletion of the tail results in σ1 trimers that oligomerize, but that are unable to encapsidate on the viral particle. Interactions between monomers of σ1 occur in the tail region, thus altering tail residues could influence the formation of the correct conformer of σ1 trimers. Such a change could affect its interaction with λ2 and encapsidation on the particle (29, 51–53). It is also possible that specific changes at residue 22 in the tail are important for the interaction of σ1 with λ2, thus either preventing interaction or weakening this interaction such that σ1 trimers do not remain stably associated with λ2. Our results add to this discovery by demonstrating that mutations in the body and head domain also reduce encapsidation of σ1. It is not expected that altering the body or head would affect encapsidation to this extent. We hypothesize that a decrease in encapsidation from σ1 mutants is a result of an altered conformation of σ1 that does not interact properly with λ2. We have shown that these changes in σ1 do not alter intracellular accumulation of the protein demonstrating that mutations do not render the protein unstable and that the effects on encapsidation with σ1 mutations arise from an assembly defect. It is also possible that σ1 that is already encapsidated is more readily ejected due to these changes. Our current work does not distinguish between these possibilities. Though σ1 reassortant viruses have been previously used to study tropism and defining receptors engaged by different reovirus serotypes, a majority of these studies have not quantified σ1 encapsidation levels or level and type of variation in the number of trimers incorporated onto particles (35, 39, 54–56). Thus, it remains possible that some of the observed phenotypes, especially those measured in vivo, are confounded by the impact of reassortment on the encapsidation efficiency of σ1. Depending on the expression level of receptors, σ1 encapsidation could impact the initial interaction between particles and the host cell.

Previous work from our lab has shown that altering σ1 affects the stability of ISVPs and decreases the encapsidation of σ1 (36). Here we show additional examples to support this idea. σ1 is not visible in cryoEM structures of virions likely because of its flexibility (51). The λ2 pentamer is in a closed conformation around σ1 in virions and ISVPs. In contrast, the λ2 pentamer is an open conformation in cores (51). This open conformation is thought to allow for the release of σ1. The structure of the ISVP* is not known but biochemical changes accompanying ISVP-to-ISVP* point to dramatic conformational changes all over the particle. During this stage, μ1 associated with the particle becomes sensitive to proteases and exposes hydrophobic residues (27, 33, 37). μ1 peptides from the N- and C-terminus of the protein are ejected along with σ1 (25, 26). Further, the particle becomes capable of transcription when supplied with NTPs (27). Based on the release of σ1 and its transcriptional activity, it is assumed that the structure of λ2 on ISVP* resembles the open structure on cores. The correlation of ISVP stability and encapsidation of σ1 may be from the fact that λ2 turrets lacking σ1 have already altered conformation to resemble ISVP*s, rendering the rest of the particle more prone to conversion to ISVP*s

ISVP stability can also be affected by σ1 via a different mechanism. We present examples where changes in σ1 produce minimal changes in encapsidation but still affect ISVPs stability. For example, T3D^F^/T3D^C^S1 and T3D^F^/T3D^C^S1-T132A both have a low level of encapsidated σ1 but T3D^F^/T3D^C^S1-T132A is more stable (Fig. 7). The mechanism behind these observations remains unknown. Though it is not known if the variety of changes in the particle accompanying ISVP-to-ISVP* transition occur concurrently or in order, our experiments with particles having a lower level of σ1 encapsidation suggest that σ1-λ2 interaction serves as a lock that maintains the stability of ISVPs and prevents premature conversion to ISVP* until an appropriate condition has been met. We think that changes in σ1 that alter its conformation can qualitatively alter its interaction with λ2 and render the particle more or less prone to ISVP* conversion. Another possibility, distinct from the σ1 - λ2 lock idea relates to the interconnectedness of reovirus capsid proteins. Though the only known interaction of σ1 in an ISVP is with λ2, λ2 in turn makes interactions with μ1 in the outer capsid and λ1 and σ2 within the core. It is possible that qualitative changes in σ1-λ2 interaction affect the conformation of λ2 in a way that its interaction with other capsid proteins is altered. This type of “ripple effect” could alter the stability of the ISVP. Indeed, our own previous work has demonstrated examples of a ripple effect where changes to one capsid protein has effects on the functions of proteins it does not interact with (36, 57). For example, changes to μ1 affect the capacity of σ1 to bind host cell receptors (57). Thus, our work highlights intricate interactions that govern the functions of the reovirus capsid.

An important question that we have not directly addressed here is the physiological relevance of having particles with lower σ1. Since σ1 serves as the primary attachment factor for reovirus, particles with lower σ1 likely cannot interact with cellular receptors and would display lower infectivity (18). However, this relationship is not linear. Previous studies indicate that the presence of 3 σ1 trimers is sufficient for particles to maintain infectivity (18). Though this was not directly tested in previous studies or our own work, presumably this number would vary with cell types due to differences in the expression level of σ1 receptors. Beyond that minimum threshold, a lower number of σ1 also appears to facilitate more rapid disassembly of the particle in the cell and a more rapid onset of viral transcription (17, 58). While ISVP* formation was not directly measured in these studies, we suggest that these data are indicative of a higher rate of ISVP-to-ISVP* formation in cells. Thus, it appears that a lower σ1 level may be advantageous at least in the environment of cultured cells to allow the virus to launch infection prior to when host responses kick in. We have also recently shown that viruses that undergo ISVP-to-ISVP* more readily elicit a lower innate immune response (59). Thus, particles with fewer σ1 or those with suboptimal σ1-λ2 interactions that promote ISVP* formation could limit the activation of the innate immune response and be beneficial to the virus. We envision however that more rapid ISVP-to-ISVP* conversion is only beneficial in an environment where ISVP formation can be controlled and only occurs in specific compartments of the cells proximal to the membrane. In other environments, such as the intestinal lumen, where ISVP formation occurs extracellularly, premature ISVP* conversion could release σ1 and μ1 peptides prior to contact with the cell and render the particle non-infectious.

## MATERIALS AND METHODS

### Cells and viruses

Murine L929 (L) cells were grown at 37°C in Joklik’s minimal essential medium (Lonza) supplemented with 5% fetal bovine serum (Life Technologies), 2 mM L-glutamine (Invitrogen). All viruses used in this study were either derived from reovirus type 3 Dearing-Field strain (T3D^F^) or type 1 Lang and were generated by plasmid-based reverse genetics (60, 61). Mutations within the T3DC and T3D^F^ S1 genes were generated by QuikChange site-directed mutagenesis (Agilent Technologies). T132A in S1 was made using the following primer pair: forward, 5’ - CGTTGCGAGAGTGGATGCCGCAGAACGTAACATTG - 3’ and reverse, 5’-CAATGTTACGTTCTGCGGCATCCACTCTCGCAACG - 3’. Chimeric σ1 viruses were obtained from the Dermody lab (University of Pittsburgh) and have been previously described (39).

### Plaque assay

Virus samples were diluted in PBS. L cell monolayers in 6-well plates (Greiner Bio-One) were infected with 100 μl of diluted virus for 1 h at room temperature. Following the viral attachment incubation, the monolayers were overlaid with 4 ml of serum-free medium 199 (Sigma-Aldrich) supplemented with 1% Bacto agar (BD Biosciences), 10 μg/ml TLCK-treated chymotrypsin (Worthington Biochemical), 2 mM L-glutamine (Invitrogen), 100 U/ml penicillin (Invitrogen), 100 μg/ml streptomycin (Invitrogen), and 25 ng/ml amphotericin B (Sigma-Aldrich). The infected cells were incubated at 37°C, and plaques were counted at 5 days post infection.

### Purification of viruses

Purified reovirus virions were generated using second- or third-passage L-cell lysate stocks of reovirus. Viral particles were Vertrel-XF (Dupont) extracted from infected cell lysates, layered onto 1.2- to 1.4-g/cm^3^ CsCl gradients, and centrifuged at 187,183 × *g* for 4 h. Bands corresponding to virions (1.36 g/cm^3^) were collected and dialyzed in virion storage buffer (150 mM NaCl, 15 mM MgCl2, 10 mM Tris-HCl [pH 7.4])(62) The concentration of reovirus virions in purified preparations was determined from an equivalence of one unit of optical density at 260 nm being 2.1 × 10^12^ virions/ml (63). Virus titer was determined by plaque assay on spinner-adapted L929 cells. At least three preparations of each viral strain were used for this study.

### Generation and analysis of ISVP-to-ISVP* conversion

ISVPs were generated *in vitro* by incubation of 2 × 10^12^ virions with 200 μg/ml of *N*α-*p*-tosyl-l-lysine chloromethyl ketone (TLCK)-treated chymotrypsin at 32°C in virion storage buffer (150 mM NaCl, 15 mM MgCl_2_, 10 mM Tris-HCl [pH 7.4]) for 20 min. ISVPs (2 × 10^12^ particles/ml) of the indicated viral strains were divided into aliquots of equivalent volumes and heated at a range of temperatures 5 min. Proteolysis was terminated by addition of 2 mM phenylmethylsulfonyl fluoride (PMSF) and incubation of reactions on ice. The reaction mixtures were cooled on ice and then digested with 0.10 mg/ml trypsin (Sigma-Aldrich) for 30 min on ice. Following addition of the SDS-PAGE loading dye, the samples were subjected to SDS-PAGE analysis. For analysis by loss in titer assays, the heated samples were used to initiate infection of L929 cells and then a plaque assay was performed.

### Selection, isolation, and sequencing of heat resistant viruses

Purified T3D^F^/T3D^C^S1 was converted to ISVPs and heated at 44°C for five minutes. Revertant viruses, which produced plaques following heat treatment were selected by plaque purification. Plaques were amplified in L-cells to obtain viral lysate stocks. To sequence gene segments in revertant viruses, viral RNA from the selected revertants was isolated by lysing infected cells with TRI Reagent (Molecular Research Center). The extracted RNA was subjected to RT-PCR using S1, M2, and L2 gene segment-specific primers. PCR products were resolved on Tris-acetate-EDTA agarose gels, purified using the QIAquick gel extraction kit (Qiagen), and sequenced using primers that cover the entire length of the S1, M2, or L2 open reading frame. Primer sequences are available upon request. To verify that the heat resistant phenotype was due to the mutation identified in sequencing, the revertant site was introduced back into T3D^F^/T3D^C^S1 and T3D^F^. These viruses were purified as described above and were analyzed for defects in plaque size, thermal inactivation, ISVP-to-ISVP* conversion, and hemagglutination.

### FLISA assay

High-affinity-binding polystyrene plates (Pierce) were coated at 4°C overnight with 1 × 10^11^ virus of purified virus diluted in 0.1 M carbonate-bicarbonate buffer at pH 9.5. Plates were blocked at 4°C for 1 h with 2.5% BSA in virion storage buffer, followed by two washes with wash buffer (0.1% BSA, 0.05% Tween, virion storage buffer). The plates were then stained at room temperature for 1 h with reovirus-specific rabbit polyclonal antiserum (1:5,000) followed by Alexa Fluor 750-labeled anti-rabbit antibody (1: 1,000) or 10 μg/ml of 9BG5 (for viruses containing T3D σ1 head) or 5C6 (for viruses containing T1L σ1 head) followed by Alexa Fluor 750-labeled anti-mouse antibody. The plate was scanned using Odyssey infrared imager (LI-COR)

### Western blot analysis

L cells were infected with T3D^F^, T3D^F^/T3D^C^S1, T3D^F^/S1-22^C^, and T3D^F^/S1-408^C^ at an MOI of 5 and the lysates were resolved on 10% SDS-PAGE gels and transferred to nitrocellulose membranes. The membranes were blocked with Starting block blocking buffer (ThermoFisher) at room temperature for 1 h followed by incubation with T3D σ1 head specific antibody (1:1000) at 4°C overnight. The membranes were washed with TBS supplemented with 0.1% Tween 20 (TBS-T) twice for 15 min and then incubated with Alexa Fluor-conjugated anti-rabbit IgG (for T3D σ1 head specific antibody) in blocking buffer. Membranes were then incubated with rabbit anti-β-actin (1:10,000) at room temperature for 1 h in blocking buffer. Membranes were then incubated in Alexa Fluor-conjugated anti-rabbit IgG (β-actin) at room temperature for 1 h. Following three washes, membranes were scanned using an Odyssey infrared imager (LI-COR). σ1/ β-actin ratios were calculated from the intensities of the σ1 and β-actin bands using Image Studio Lite software (Li-COR).

### Agarose gel separation of reovirus particles by σ1 content

A total of 1 × 10^11^ virus particles were resuspended in dialysis buffer, mixed with 2× agarose gel loading dye (NEB), and resolved on 1% ultrapure agarose gel (Invitrogen) in 1 × TAE, pH 7.2, at a constant voltage of 25 V for 18 h. The gel was stained with a Novex colloidal blue staining kit (Invitrogen) for 6 h and destained overnight in water. The gel was scanned using an Odyssey infrared imager (LI-COR). The intensity of each band, representing particles with a specific number of σ1 trimers, was quantified using Image Studio Lite software. The data were represented as the percentage of that species within the virus preparation.

### Measurement of stability of ISVPs with different amounts of σ1

A total of 1 × 10^11^ virus particles were resuspended in dialysis buffer, mixed with 2× agarose gel loading dye (NEB), and resolved on 1% ultrapure agarose gel (Invitrogen) in 1 × TAE, pH 7.2, at a constant voltage of 25 V for 18 h. The same sample was run in 8 lanes in parallel. One lane was cut out and stained in ethidium bromide for 3 hours. Once bands were visible, the remaining gel was sectioned horizontally into three groups (high, medium, and low encapsidation). 1 mL of virion storage buffer was added before sonication. Samples were then nutated for at least 3 days at 4 degrees. Samples were then converted to ISVPs by chymotrypsin treatment and heated at various temperatures. Heated ISVPs were then used to infect plaque assays to measure change in titer at different temperatures.

### Statistical analysis

The reported values represent the mean of three independent, biological replicates. Error bars indicate standard deviation. P values were calculated using Student’s t test (two-tailed, unequal variance assumed), One-way ANOVA, or Two-way ANOVA, Dunnett’s with multiple comparisons.

